# Crosstalk of growth factor receptors at plasma membrane clathrin-coated sites

**DOI:** 10.1101/2024.05.16.594559

**Authors:** Marco A. Alfonzo-Méndez, Marie-Paule Strub, Justin W. Taraska

## Abstract

Cellular communication is regulated at the plasma membrane by the interactions of receptor, adhesion, signaling, exocytic, and endocytic proteins. Yet, the composition and control of these nanoscale complexes in response to external cues remain unclear. Here, we use high-resolution and high-throughput fluorescence imaging to map the localization of growth factor receptors and related proteins at single clathrin-coated structures across the plasma membrane of human squamous HSC3 cells. We find distinct protein signatures between control cells and cells stimulated with ligands. Clathrin sites at the plasma membrane are preloaded with some receptors but not others. Stimulation with epidermal growth factor induces a capture and concentration of epidermal growth factor-, fibroblast growth factor-, and low-density lipoprotein-receptors (EGFR, FGFR, and LDLR). Regulatory proteins including ubiquitin ligase Cbl, the scaffold Grb2, and the mechanoenzyme dynamin2 are also recruited. Disrupting FGFR or EGFR individually with drugs prevents the recruitment of both EGFR and FGFR. Our data reveals novel crosstalk between multiple unrelated receptors and regulatory factors at clathrin-coated sites in response to stimulation by a single growth factor, EGF. This behavior integrates growth factor signaling and allows for complex responses to extracellular cues and drugs at the plasma membrane of human cells.

**Significance:** Classically, receptor pathways including epidermal growth factor receptor and fibroblast growth factor receptor were thought of as independent systems. Yet, the plasma membrane is a complex environment where proteins interact, cluster, signal, and associate with organelles. For example, after EGF activation, EGFR is captured at sites on the inner plasma membrane coated with the protein clathrin. This causes clathrin to grow flat across the adherent membrane. Here, we observe co-capture along with EGFR of the related receptor FGFR and unrelated LDLR by clathrin after EGF stimulation. This is specific as other receptors are unaffected. Thus, separate but specific receptor systems co-assemble and signal to each other at nanoscale zones on the plasma membrane organized by clathrin. This provides new avenues for treating diseases like cancer.

## Introduction

Receptor tyrosine kinases (RTKs) are key plasma membrane (PM) receptors in humans. RTKs are activated by extracellular growth factors including epidermal growth factor (EGF) and fibroblast growth factor (FGF). RTKs regulate cell proliferation, differentiation, survival, migration, and development (1). RTK dysfunction leads to uncontrolled cell growth and cancer (2). For this reason, commonly used chemotherapies target RTKs to prevent their activation (3). However, these drugs can cause adverse side effects and cell resistance. Hence, understanding how RTKs work is key to designing better anti-cancer treatments.

One of the most well studied RTKs is the Epidermal Growth Factor Receptor (EGFR). EGFR is a single-pass transmembrane protein with an extracellular N-terminal EGF-binding domain and an intracellular C-terminal with a tyrosine kinase domain (4). EGF binding induces receptor dimerization and cross-tyrosine phosphorylation (5). In turn, the added phosphates provide docking sites for multiple proteins to activate signaling cascades at the PM (6, 7).

Aside from signaling-associated proteins (8), EGFR function is modulated by clathrin-mediated endocytosis —the major receptor internalization pathway in humans (9). EGF triggers changes in the structure of a long-lived subset of coats known as flat clathrin lattices (FCLs). Over minutes, FCLs grow and multiply, coating the inner adherent PM to organize signaling. The assembly and growth of FCLs is regulated by the co-capture of EGFR, the adhesion receptor β5-integrin, and the tyrosine kinase Src (10-13). This tri-partite axis enhances the activity of EGFR and integrates two different systems—growth factor signaling and cell adhesion—at nanoscale molecular hubs. However, whether FCLs facilitate the physical integration and activation of other signaling pathways across the PM remains unknown.

While receptors have been mostly studied in isolation, it is becoming clear that different members of the RTK family can crosstalk with other receptors (14-16). For example, EGFR inhibitors were identified in a screen for drugs that improve outcomes in cancers driven by the fibroblast growth factor receptor (FGFR) (17). EGFR and FGFR are both RTKs, but they are not thought to oligomerize on the PM and they bind to and are active by different ligands —EGF and FGF— (18, 19). How this crosstalk happens is unclear. Is it a physical interaction, one that occurs through dynamic signaling, or one that occurs through scaffolding at the plasma membrane?

Here, we explore if stimulation with EGF or FGF increases the proximity or co-clustering of different receptors at signaling domains at the plasma membrane organized by clathrin-coated structures (CCSs). We find that EGF causes the rapid co-capture of the related EGFR and FGFR receptors and the unrelated low density lipoprotein receptor into clathrin-coated sites at the ventral PM. Other receptors including G-protein coupled receptors are not affected after EGF stimulation. Specific signaling and endocytic proteins also are co-captured after EGF stimulation included the ubiquitin ligase Cbl, endocytic proteins including Eps15/R and dynamin2, and the EGFR-binding scaffold protein Grb2. Drugs that individually target a single RTK blocked the co-clustering of both FGFR and EGFR at these sites. We conclude that different receptor systems interact at the nanoscale in clathrin coated sites and activation of one can induce capture of the other. This crosstalk links multiple signaling systems both inside and outside clathrin coated sites, providing new opportunities to perturb and control these receptors in both health and disease.

## Results

### Quantitative measurements of EGFR recruitment into CCSs

We have previously shown that EGFR activation by EGF induces the growth of flat clathrin lattices (FCLs). In turn, the clustering of EGFR in FCLs enhances EGFR signaling at the PM (Fig. 1*A*). Thus, FCLs are signaling hubs that can dynamically harbor signaling proteins such as the tyrosine kinase Src and the cell adhesion receptor β5-integrin (10). But are these EGF-induced signaling hubs specific for the EGFR/Src/β5-integrin axis? To address this question, we used an automated two-color image correlation pipeline and measured the presence of signaling and endocytic adaptor proteins across thousands of individual CCSs (20). In Figure 1, we show EGFR as an example of our pipeline to track the nanoscale dynamics of proteins before and after EGF stimulation.

**Fig. 1.**
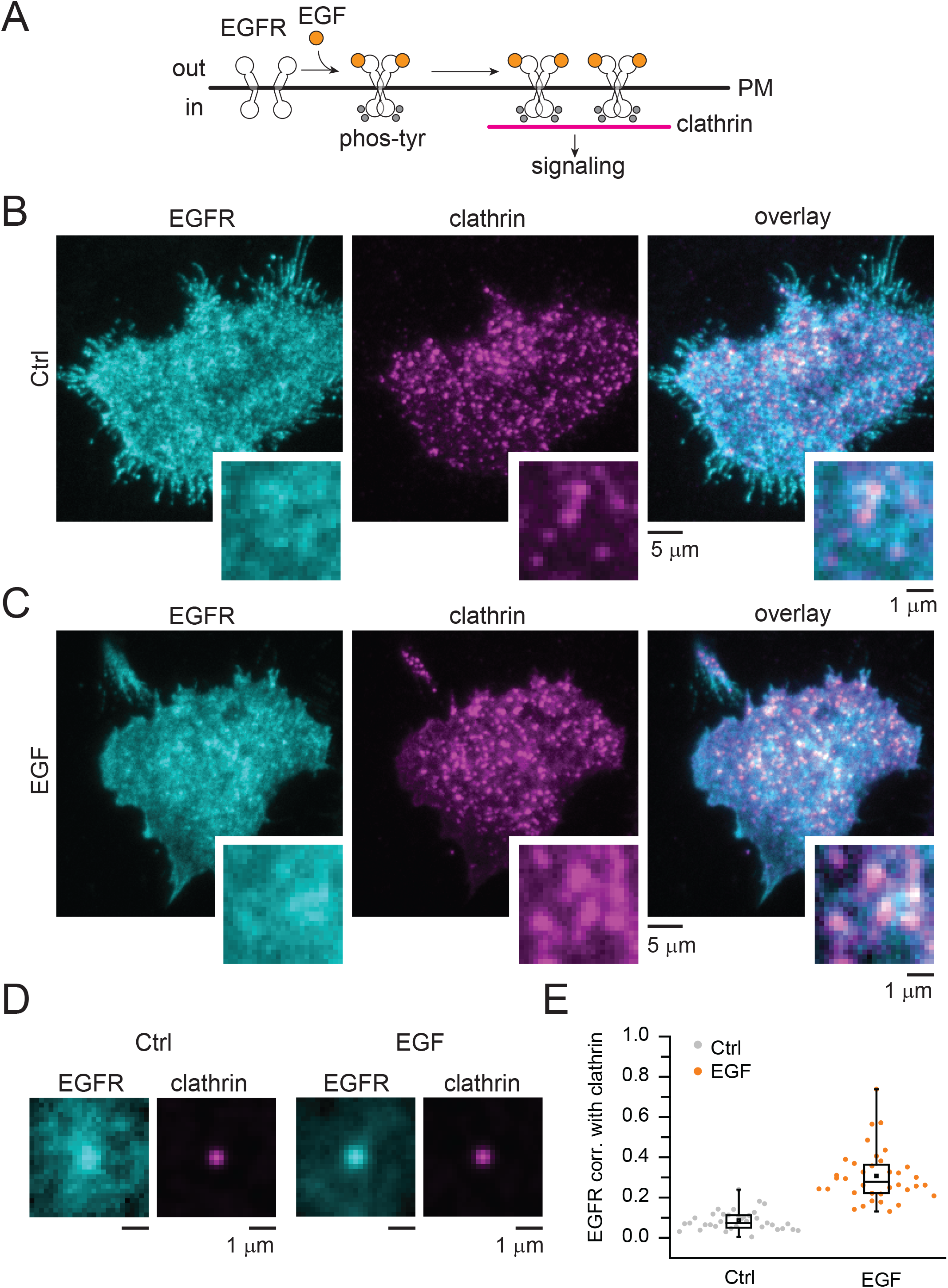
Quantitative measurements of EGFR recruitment into CCSs. (*A*) Cartoon depicting EGFR dimerization, phosphorylation at tyrosine residues (grey circles), and recruitment into clathrin after EGF binding (orange circles). (*B*-*C*) Representative two-color TIRF images of genome edited HSC3 expressing EGFR-GFP (shown in cyan) and transfected with mScarlet-CLCa (shown in magenta) before (Ctrl) or after 50 ng/mL EGF stimulation for 15 min. Scale bar is 5 μm; insets scale bar is 1 μm. (*D*) Representative images obtained after averaging small regions extracted from the magenta channel, along with the corresponding region of the cell from the cyan channel. All regions are normalized to the brightest pixel. Scale bar is 1 μm. (*E*) Automated correlation analysis between clathrin and EGFR. Dot box plots show median extended from 25th to 75th percentiles, mean (square), and minimum and maximum data point whiskers with a coefficient value of 1.5. N = 3 biologically independent experiments. EGFR epidermal growth factor receptor, CCSs clathrin-coated structures, PM plasma membrane, TIRF total internal reflection fluorescence, CLCa clathrin light chain a, EGF epidermal growth factor.

Figure 1*B* shows total internal reflection fluorescence (TIRF) microscope images of human HSC3 squamous cells engineered to express EGFR-GFP (cyan) at native levels. HSC3 cells have been used as a model to study EGFR endocytosis and human head and neck carcinoma (21). These cells were transfected with clathrin light chain fused to the red fluorescent protein mScarlet (magenta) to mark single clathrin-coated structures at the bottom PM. In control cells (Ctrl), EGFR showed little overlap with clathrin (Fig. 1*B*). In contrast, the EGFR-clathrin overlap increased after EGF stimulation, shown as pseudo-colored white in the overlay images (Fig. 1*C*, right panel). To measure these changes, we used TIRF clathrin images as targets to extract small regions of the cell in both protein-specific color channels. When thousands of single clathrin light chain-mScarlet images are normalized and averaged, we observed a sub-diffraction centered bright spot. In control cells, EGFR-GFP appeared as a diffuse signal across the average image (Fig. 1*D*, left panels). After EGF simulation, the average EGFR signal is enhanced in the center (Fig. 1*D*, right panels). At each structure we used a Pearson’s correlation function to measure the degree of overlap and changes in correlation between the target protein and clathrin in a 3-pixel radius area (501 nm) surrounding the clathrin site. A value of 1 is maximum correlation and a value of -1 is maximum anti-correlation according to the function. Figure 1*E* shows that across many structures, cells, and replicates, the correlation between EGFR and clathrin is enhanced after EGF stimulation (Ctrl. 4263 structures, EGF. 3580, 36 cells, p-value = 2.39E-14). The correlation values with clathrin in control cells are 0.09±0.008 SEM and 0.31±0.02 SEM in EGF stimulated cells. We also observed consistent changes when using larger areas of analysis (12-pixel radius), indicating that the direction of changes was insensitive to the specific analysis parameters. Overall, our data demonstrates the robustness of our method to assess the nanoscale association and changes in association of proteins at clathrin-coated structures during EGF signaling.

### Distinct receptors differentially locate in CCSs in response to EGF

EGFR is captured by and causes changes to the structure of clathrin sites visible by electron microscopy (10). However, along with EGFR, the plasma membrane is populated by a wide array of receptors and signaling proteins. We hypothesized that other proteins might re-distribute with respect to clathrin after EGF simulation. To address this, we performed a screen of 53 different candidate proteins, and we evaluated changes in their correlation with clathrin (SI Appendix, Fig. 1). From this screening, we chose 7 well-studied receptors that are related or unrelated in sequence, structure, or function to RTKs. We included the EGFR-related fibroblast growth factor receptor (FGFR) and HER2, the unrelated low density lipoprotein receptor (LDLR) and transferrin receptor (TfR), and a set of G-protein coupled receptors including the lysophosphatidic receptor 1 (LPAR1), β_2_-adrenergic receptor (β_2_-AR), and the α_1B_-adrenergic receptor (α_1B_-AR) (Fig. 2*A*). In these experiments, HSC3 cells were co-transfected with a single fluorescently labeled receptor and fluorescent clathrin light chain, stimulated with EGF, fixed, and imaged with TIRF. Figure 2*B* shows representative overlay images of cells expressing the receptors tested together with clathrin. Figure 2*C* shows quantitation of the correlation between clathrin and the different receptors across multiple biological replicates, cells, and thousands of structures. Surprisingly, two receptors that do not bind to EGF ligand showed increased correlation with clathrin after EGF stimulation (Fig. 2*C*).

**Fig. 2.**
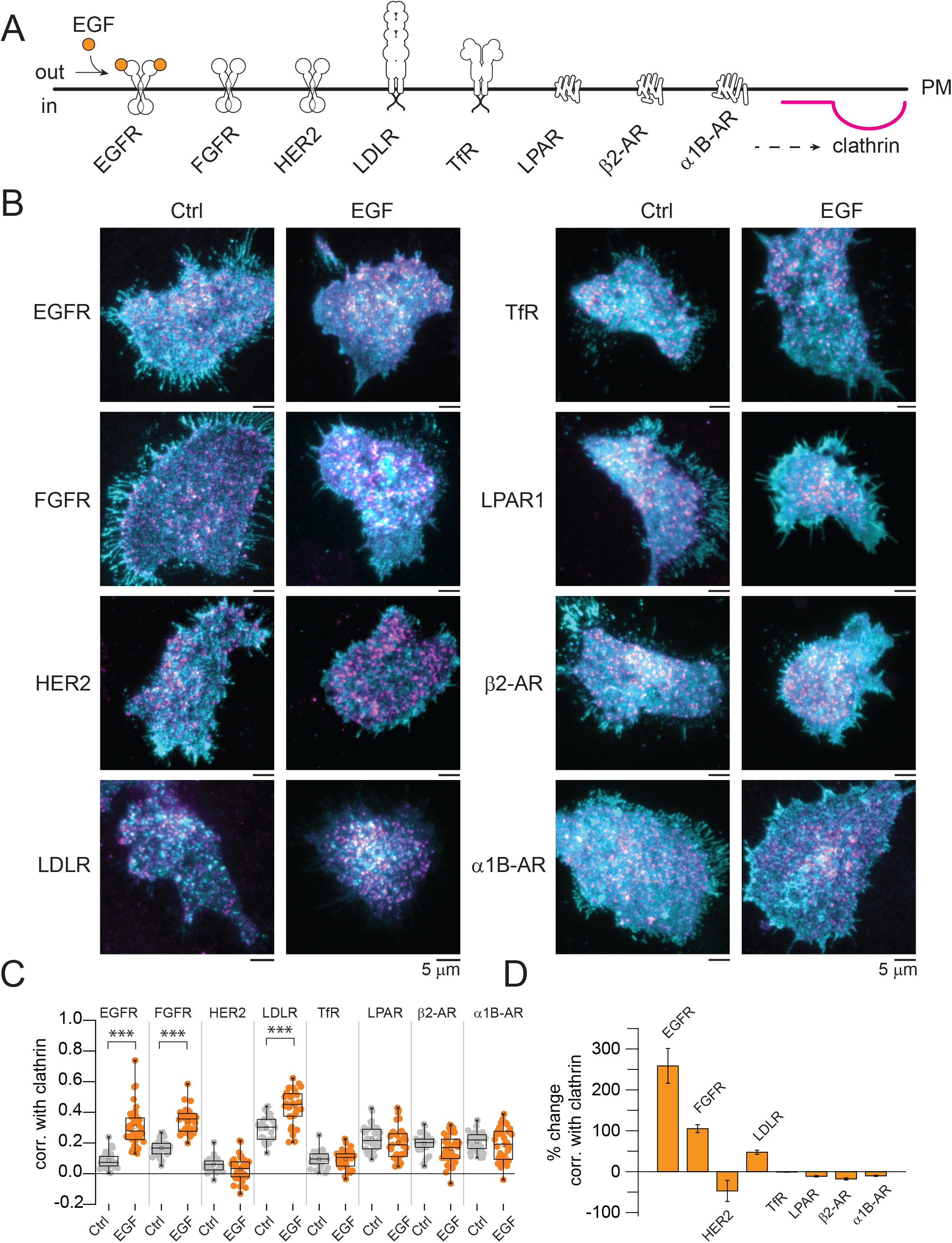
Distinct receptors differentially locate in CCSs in response to EGF. (*A*) Cartoon depicting the different receptors whose localization was evaluated upon EGF stimulation (orange circles). (*B*) Representative two-color TIRF images of HSC3 transfected with the indicated receptors and mScarlet-CLCa before (Ctrl) or after 50 ng/mL EGF stimulation for 15 min. Scale bar is 5 μm. (*C*) Automated correlation analysis between clathrin and the indicated receptor. Dot box plots show median extended from 25th to 75th percentiles, mean (square), and minimum and maximum data point whiskers with a coefficient value of 1.5. Significance was tested by a two-tailed t-test. (*D*) Percent change of correlation between clathrin and the indicated receptor. N = 3 biologically independent experiments. CCSs clathrin-coated structures, EGF, epidermal growth factor, EGFR epidermal growth factor receptor, TIRF total internal reflection fluorescence, CLCa clathrin light chain a, FGFR fibroblast growth factor, Her2 human epidermal growth factor receptor 2, LDLR low-density lipoprotein receptor, TfR transferrin receptor, LPAR1 lysophosphatidic acid receptor 1, β_2_-AR β2 adrenergic receptor, α_1B_-AR a α1B-adrenergic receptor.

Specifically, the related FGFR, which normally binds to FGF, was recruited into CCSs. The unrelated LDLR, which binds to and internalizes low density lipoproteins, also associated with CCSs after EGF stimulation (Fig. 2*C*). In contrast, the HER2 receptor, known to associate with EGFR (22), was not redistributed to clathrin sites (Fig. 2*C*). The correlation with clathrin of the other receptors (TfR, LPAR, β_2_-AR, and α_1B_-AR), showed no changes or even slight decreases as compared to control cells. To quantitate these changes, we calculated the percent change in correlation between clathrin and each receptor. EGFR showed the largest response to EGF (259%), followed by FGFR (105%), and then LDLR (48%) (Fig. 2*D*). These results showed that CCSs at the PM not only capture EGFR but other receptors such as FGFR and LDLR after EGF stimulation. Hence, clathrin lattices are more global signaling hubs than previously proposed.

### Distinct receptors are captured into CCSs in response to their corresponding agonist

While we saw no changes in G-protein coupled receptors after EGF stimulation, past studies showed that these receptors are captured by clathrin after stimulation with their corresponding native ligands (22-27). Thus, as a control, we determined if a subset of receptors associate with clathrin when stimulated with their native ligands. Figure 3 shows that LPAR clusters into clathrin when exposed to its natural ligand lysophosphatidic acid (LPA), and similar to past work (23), β2-AR clusters with clathrin after isoproterenol (Iso) stimulation (Fig. 3). In turn, FGFR is also captured by clathrin after stimulation with its natural ligand, FGF. We did not detect an increased association of α_1B_-AR with clathrin after stimulation with noradrenaline (NA). These data indicate that different receptors are recruited into clathrin after stimulation with their native specific ligands. Yet, only FGFR, EGFR, and LDLR are stimulated to cluster with clathrin after EGF stimulation.

**Fig. 3.**
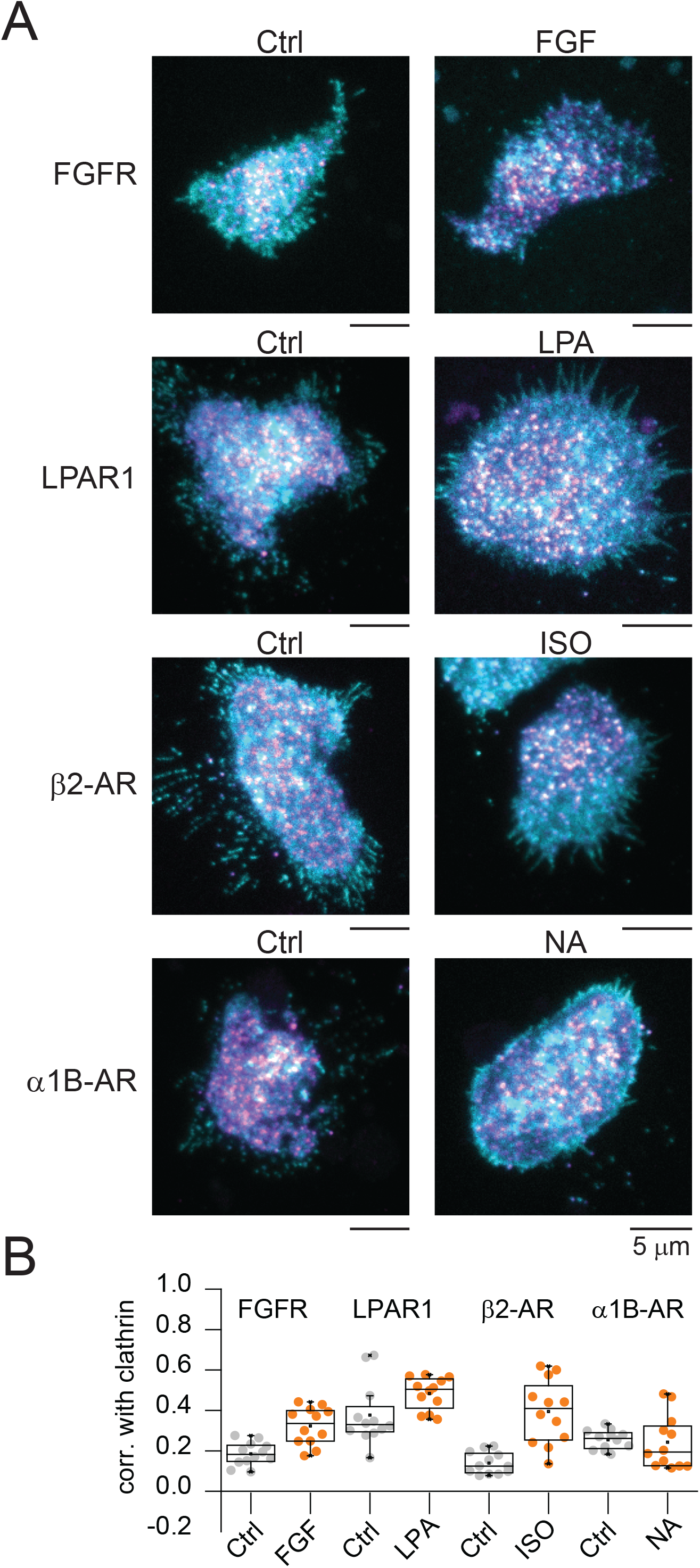
Receptors are captured into CCSs in response to their corresponding agonist. (*A*) Representative two-color TIRF images of HSC3 transfected with the indicated receptors and mScarlet-CLCa before (Ctrl) or after stimulation for 15 min with their corresponding agonists (50 ng/mL FGF, 10 μM lysophosphatidic acid, 1 μM isoproterenol, and 10 μM noradrenaline). Scale bar is 5 μm. (*B*) Automated correlation analysis between clathrin and the indicated receptor. Dot box plots show median extended from 25th to 75th percentiles, mean (square), and minimum and maximum data point whiskers with a coefficient value of 1.5. N = 3 biologically independent experiments. CCSs clathrin-coated structures, TIRF total internal reflection fluorescence, CLCa clathrin light chain a, FGFR, fibroblast growth factor receptor, FGF fibroblast growth factor, LPAR1 lysophosphatidic acid receptor 1, LPA lysophosphatidic acid, β_2_-AR β_2_-adrenergic receptor, ISO isoproterenol, α_1B_-AR a α_1B_-adrenergic receptor, NA noradrenaline.

### Distinct endocytic proteins differentially locate in CCSs in response to EGF

Multiple endocytic and signaling adaptors are known to be biochemically connected to EGFR in different ways (28-31).

However, changes in their location at the PM during growth factor signaling is unknown. Next, we tested the response of a set of endocytic and signaling proteins. Our hypothesis was that these proteins would show a corresponding co-capture with clathrin to help recruit receptors such as FGFR and LDLR into clathrin sites during EGF stimulation. Figure 4 shows analysis for 10 regulatory proteins before and after EGF stimulation. We observed a diversity of behaviors. Endocytic adaptors like intersectin1 (ITSN1), β-arrestin2 (β-arr2), and Dab2, showed no changes or decreases in their correlation with clathrin. Whereas Eps15, Eps15R, ARH and NUMB showed mild increases. However, ARH, Dab2, Esp15, Eps15R, and ITSN1 are pre-associated with clathrin before stimulation (Figs. 4). Of note, three proteins showed substantial increases with clathrin: 1) the mechanoenzyme and actin bundling protein dynamin2; 2) the ubiquitin ligase Cbl; 3) and the scaffold protein Grb2 (Fig. 4C). Thus, EGF triggers changes in the location of specific endocytic and regulatory proteins forming CCSs.

**Fig. 4.**
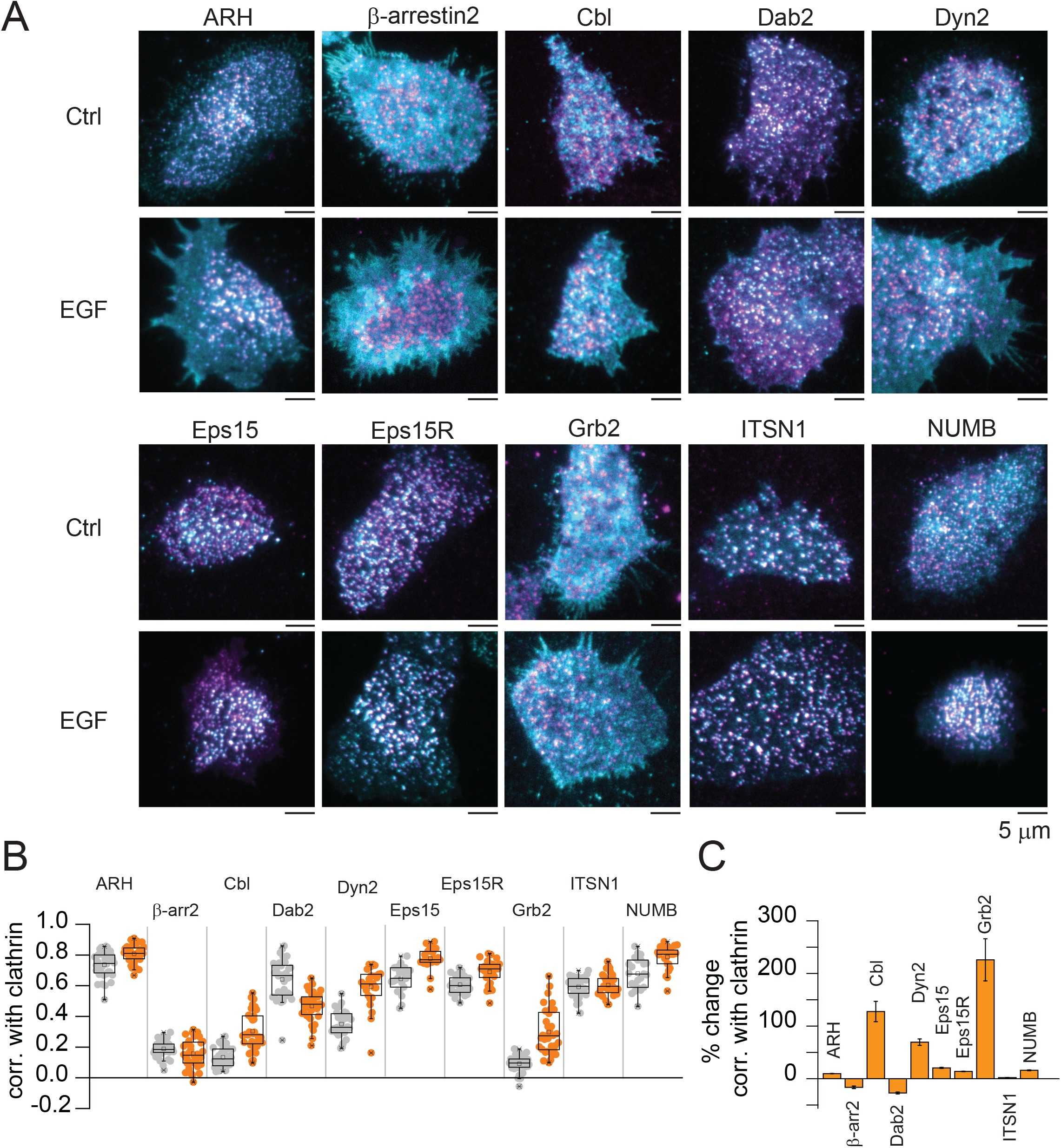
Distinct endocytic proteins differentially locate in CCSs in response to EGF. (*A*) Representative two-color TIRF images of HSC3 transfected with the indicated endocytic proteins and mScarlet-CLCa before (Ctrl) or after 50 ng/mL EGF stimulation for 15 min. Scale bar is 5 μm. (*B*) Automated correlation analysis between clathrin and the indicated endocytic protein. Dot box plots show median extended from 25th to 75th percentiles, mean (square), and minimum and maximum data point whiskers with a coefficient value of 1.5. (*C*) Percent change of correlation between clathrin and the indicated endocytic protein. N = 3 biologically independent experiments. CCSs clathrin-coated structures, EGF, epidermal growth factor, TIRF total internal reflection fluorescence, CLCa clathrin light chain a, ARH autosomal recessive hypercholesterolemia adaptor protein, β-arr2 β-arrestin 2, Cbl Casitas B-lineage lymphoma ubiquitin ligase, Dab2 Disabled homolog 2, Dyn2 dynamin2, Eps15 epidermal growth factor receptor substrate 15, Eps15R Eps15 related protein, Grb2 growth factor receptor-bound protein 2, ITSN1 intersectin 1, NUMB protein numb homolog.

### EGFR recruitment into CCSs in response to both EGF and FGF requires EGFR kinase activity

To examine the mechanisms leading to co-capture of receptors at CCS, we focused on EGFR and FGFR. Two drugs are known to selectively block these receptors. Gefitinib is a widely used anti-cancer drug that targets the kinase domain of EGFR and prevent its activity (32). PD-166866 is a competitive antagonist of the FGFR kinase domain (33). We used these two selective inhibitors to investigate how blocking the kinase activity of either EGFR or FGFR affected the capture of the receptors into CCSs.

First, we tested how blocking both EGFR and FGFR activity affected the clustering of the EGFR receptor into clathrin after stimulation with EGF and FGF (Fig. 5*A*). Figure 5*B* shows representative images of EGFR-GFP expressing cells co-transfected with clathrin light chain-mScarlet before and after a series of perturbations including incubation with EGF, EGF with the EGFR inhibitor gefitinib (EGF+Gefi), and EGF with the FGFR inhibitor PD-166866 (EGF+PD) (Fig. 5*B*). We observed that EGF induced the capture of EGFR into clathrin sites and this effect was abolished by gefitinib. Surprisingly, blocking FGFR kinase activity with PD-166866 decreased EGFR recruitment into CCSs in response to its natural ligand: EGF (Fig. 5*C*). Next, we assessed changes in the location of EGFR, but in response to FGFR natural ligand: FGF (Fig. 5*D*). Unexpectedly, we detected an increase in the correlation of EGFR after stimulation with FGF. This effect was blocked by gefitinib (FGF+Gefi), but not by PD-166866 (FGF+PD) (Fig. 5*E*). These data indicates that: 1) the EGFR recruitment into CCSs can be triggered by both its natural ligand —EGF— and by another growth factor, FGF; 2) EGFR kinase activity is needed for the clustering of EGFR induced both by EGF and FGF; and 3) FGFR kinase activity boosted EGF-induced EGFR clustering. Altogether, these results suggest that EGFR and FGFR are spatially connected by CCSs at the PM.

**Fig. 5.**
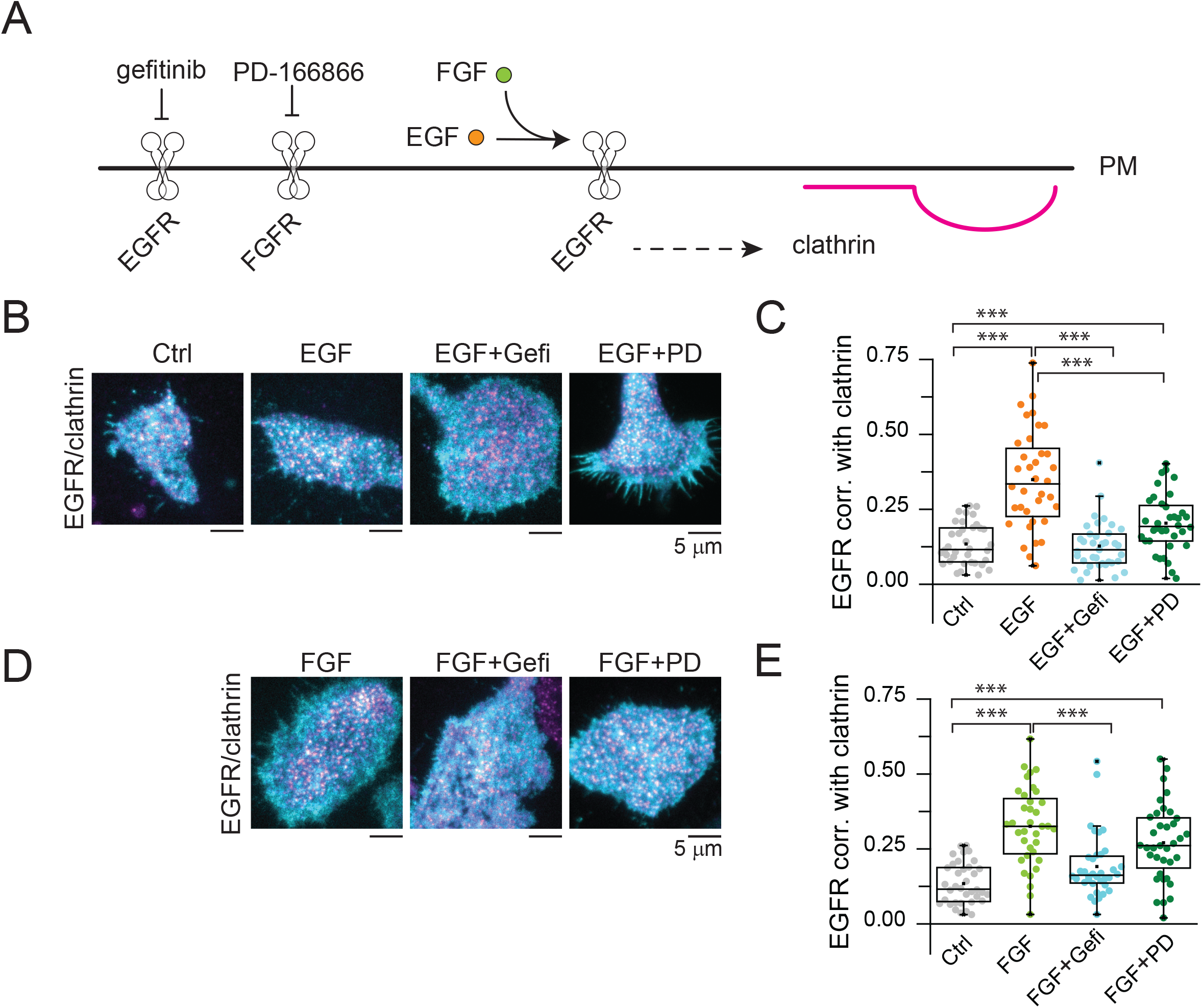
EGFR recruitment into CCSs in response to both EGF and FGF requires EGFR kinase activity. (*A*) EGFR localization in clathrin was evaluated upon EGF stimulation (orange circles) in the presence or absence of EGFR and FGFR specific inhibitors (gefitinib and PD 166866, respectively). (*B*) Representative two-color TIRF images of HSC3 transfected with EGFR and mScarlet-CLCa before (Ctrl) and after treatment with either 50 ng/mL EGF alone for 15 min, or in the presence of 10 μM gefitinib (EGF+Gefi) or 10 μM PD-166866 (EGF+PD). (*C*) Automated correlation analysis between clathrin and EGFR from cells in *B*. (*D*) Representative two-color TIRF images of HSC3 transfected with EGFR and mScarlet-CLCa before (Ctrl) and after treatment with either 50 ng/mL FGF alone for 15 min, or in the presence of 10 μM gefitinib (EGF+Gefi) or 10 μM PD-166866 (EGF+PD). Scale bar is 5 μm. (*E*) Automated correlation analysis between clathrin and EGFR from cells in *D*. Dot box plots show median extended from 25th to 75th percentiles, mean (square), and minimum and maximum data point whiskers with a coefficient value of 1.5. Significance was tested by a two-tailed t-test. N = 3 biologically independent experiments. EGFR, epidermal growth factor receptor, CCSs clathrin-coated structures, EGF epidermal growth factor, FGF fibroblast growth factor, TIRF total internal reflection fluorescence, CLCa clathrin light chain a.

### RTK blockers perturb the recruitment of FGFR into clathrin-coated sites on ligand activation

To further explore the crosstalk between EGFR and FGFR, we examined the clustering of FGFR after stimulation with EGF and the perturbations described above (Fig. 6*A*). Surprisingly, EGF caused clustering of the related RTK —FGFR— into CCSs (Fig. 6*B*). The FGFR recruitment into CCSs triggered by EGF was blocked by gefitinib (EGF+Gefi) but not by PD-166866 (EGF+PD) (Fig. 6*C*). Finally, we evaluated changes in the correlation between FGFR and clathrin, in response to the FGFR natural ligand: FGF (Fig. 6*D*). As expected, FGF caused an increase in FGFR correlation with clathrin (Fig. 6*E*). This increase was not disturbed by inhibiting the EGFR kinase activity with gefitinib (FGF+Gefi). In contrast, the FGFR inhibitor PD-166866 blocked the FGFR recruitment into CCSs induced by FGF (FGF+PD). In summary: 1) the capture of FGFR into CCSs can be induced by both its natural ligand — FGF —and by another growth factor, EGF; 2) EGFR kinase activity is needed for the clustering of FGFR induced by EGF but not by FGF; and 3) FGFR kinase activity is required for FGFR clustering triggered by FGF but not by EGF. These data further confirm the signaling crosstalk between both the EGFR and FGFR systems.

**Fig. 6.**
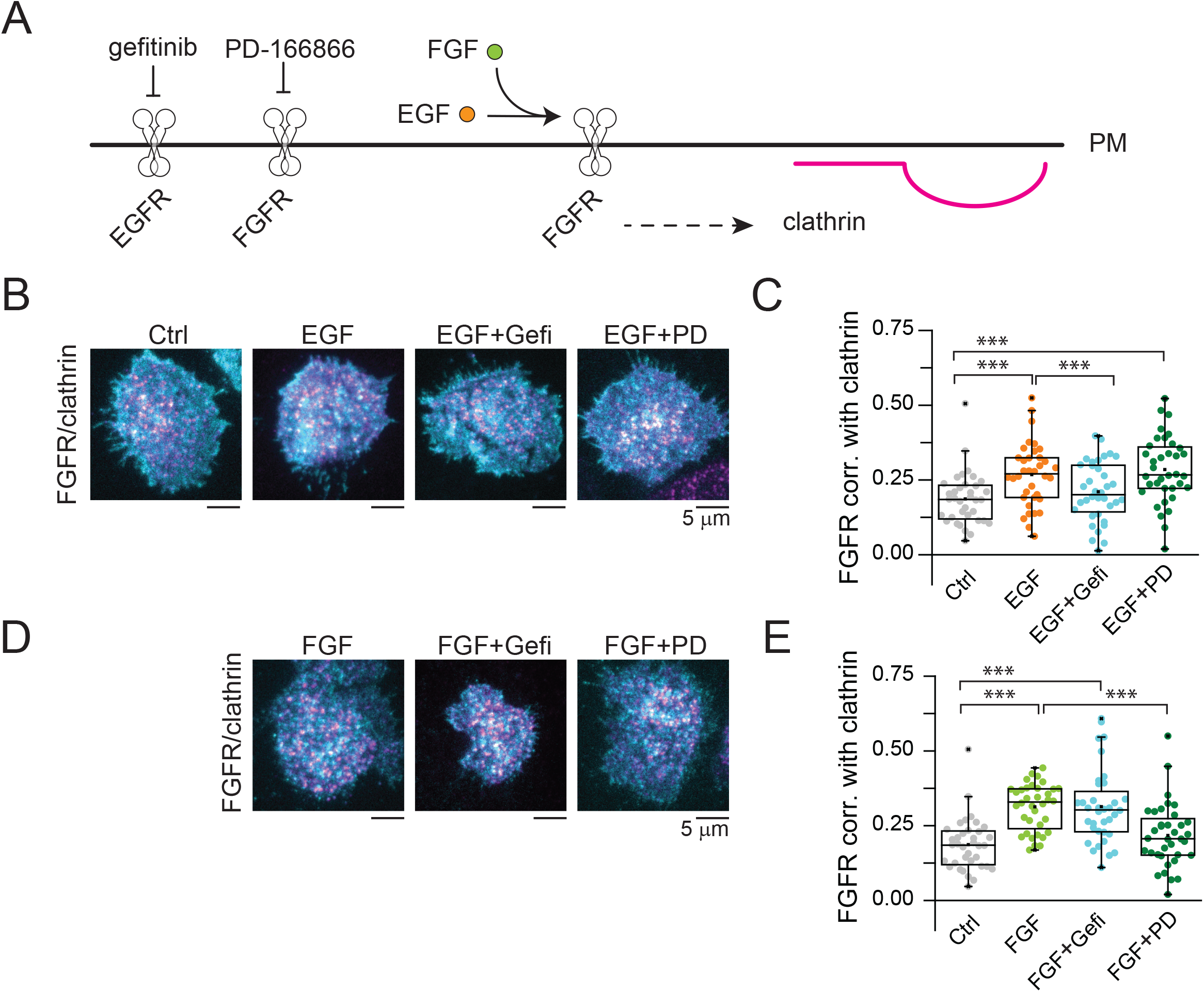
FGFR recruitment into CCSs in response to both EGF and FGF is EGFR kinase independent. (*A*) FGFR localization in clathrin was evaluated upon FGF stimulation (green circles) in the presence or absence of EGFR and FGFR specific inhibitors (gefitinib and PD 166866, respectively). (*B*) Representative two-color TIRF images of HSC3 transfected with FGFR and mScarlet-CLCa before (Ctrl) and after treatment with either 50 ng/mL EGF alone for 15 min, or in the presence of 10 μM gefitinib (FGF+Gefi) or 10 μM PD-166866 (FGF+PD). (*C*) Automated correlation analysis between clathrin and FGFR from cells in *B*. (*D*) Representative two-color TIRF images of HSC3 transfected with FGFR and mScarlet-CLCa before (Ctrl) and after treatment with either 50 ng/mL FGF alone for 15 min, or in the presence of 10 μM gefitinib (FGF+Gefi) or 10 μM PD-166866 (FGF+PD). Scale bar is 5 μm. (*E*) Automated correlation analysis between clathrin and FGFR from cells in *D*. Dot box plots show median extended from 25th to 75th percentiles, mean (square), and minimum and maximum data point whiskers with a coefficient value of 1.5. Significance was tested by a two-tailed t-test. N = 3 biologically independent experiments. EGFR, epidermal growth factor receptor, CCSs clathrin-coated structures, EGF epidermal growth factor, FGF fibroblast growth factor, TIRF total internal reflection fluorescence, CLCa clathrin light chain a.

## Discussion

The diverse signaling pathways of human cells are integrated at multiple levels. However, when, where, and how this crosstalk occurs within the complexity of the PM remains unclear. Clathrin lattices act as nanoscale signaling sites at the PM where multiple signaling proteins including EGFR, integrins, and Src are dynamically concentrated and co-regulated through phosphorylation (10, 11, 34). Indeed, signaling from some ligand-bound receptors can even occur after clathrin is taken up into the cell as a vesicle (35, 36). Here, we show that EGF induced a robust co-clustering of receptors that do not naturally bind to EGF, such as FGFR and LDLR, into CCSs along with EGFR. Similar capture of EGFR was seen after FGF activation. Along with these receptors, key endocytic, enzymatic, and scaffolding proteins are also recruited including Cbl, Grb2, dynamin2, and Eps15/R. Drugs that specifically inhibit either EGFR or FGFR disrupt receptor recruitment into CCSs. Our data reveal a crosstalk between different RTK signaling pathways that is generated by changes to clathrin-coated signaling domains at the adherent PM.

Over the last few years clathrin has been shown to be a central organizer of signaling and adhesion (37). This is a function separate from its role in generating membrane curvature and transport vesicles for endocytosis (38-40). Cells use clathrin as small adhesion sites during migration and division (41-43). Of note, in muscle cells, clathrin is used to fortify the PM at sites of actin (44). Cells also use clathrin to concentrate and organize receptors (35). Presumably, clathrin can transition from a signaling hub into an endocytic vesicle-forming site, packaging signaling or adhesion domains into vesicles for transport into the cell when they are not needed. Here, we show a collective behavior of specific receptors at clathrin-organized domains in response to growth factor stimulation.

Three different receptors were clustered into clathrin after EGF stimulation. Both EGFR and FGFR were responsive to both EGF and FGF stimulation. Surprisingly, LDLR was also co-captured by clathrin after growth factor stimulation. Why this occurs is still unclear as LDLR, FGFR, and EGFR are not thought to bind to one another and are unrelated in sequence and structure. Thus, a recruited protein likely plays a role in co-capturing these proteins or intracellular signaling systems can cross-activate each receptor independently. The capture of EGFR and FGFR is dependent on the phosphorylation of the intracellular domains of these proteins (45, 46). How LDLR is affected is less clear. There is currently conflicting evidence that LDLR is allosterically activated to bind to clathrin (47, 48). Indeed, LDLR might cluster and bind to clathrin directly without adaptors. Yet, phosphorylation of the LDLR related protein by PKCα has been shown to modulate binding to AP2 (49). Many of the receptors we imaged, including TfR and G-protein coupled receptors, were not concentrated in CCSs in response to EGF. Thus, how this differential response in receptor behavior is generated is still unclear and will require future work.

Through an unbiased screen we mapped multiple regulatory proteins at clathrin sites after EGF stimulation. Several proteins showed dynamic changes. The most prominent and strongest increases were seen with Cbl, Grb2, dynamin2, with smaller but significant increases of ARH, Eps15/R, Numb, and a surprising loss of Dab2. Many of these proteins are known binding partners of EGFR or are enzymes that act on growth factor rector systems (28-31). Likewise, many of these factors have SH3 domains that are thought to organize and crosslink complexes during signaling or endocytosis (50, 51). We have not identified a specific protein that might be the master recruiter of LDLR, EGFR, or FGFR during growth factor activation. Given that many of the receptors are ubiquitinated, a ubiquitin ligase such as Cbl is a prime candidate for generating this general response. The ubiquitinated receptors whether they be EGFR, FGFR, or LDLR would be able to co-cluster with ubiquitin binding proteins such as epsins at the growing clathrin coated sites. This could even be facilitated by a phase-like transition in these complexes (52). Yet, many G-protein-coupled receptors are also ubiquitinated (53). Thus, this ubiquitination would need to be specific to these three receptors, EGFR, FGFR, and LDLR only after EGF stimulation. More work in needed to unravel this complexity. It is interesting to note that some endocytic proteins showed decreases on stimulation. This behavior could be the result of crowding/filling of the limited binding sites on a single clathrin structure as a new set of proteins are recruited during growth factor stimulation.

What is the functional consequence of EGF and FGF actions on other receptors? Clustering of EGFR at clathrin sites enhances the signaling output of EGFR. We have not yet determined the direct output of clustering FGFR or LDLR at clathrin sites after EGF stimulation. Likewise, the functional consequence of capturing EGFR after FGF stimulation is unresolved. Additional work is needed to answer these questions. However, it is likely that similar to EGF, induced clustering of EGFR either enhances endocytosis, enhances signaling, or induces sequestration of LDLR or FGFR. On its own, our data point to a much more interconnected system of these receptors in human cells. Previous models proposed them to be fully independent with unique and non-overlapping ligands and signaling outputs. The fact that drugs against either receptor influenced the behavior of the other point to direct signaling through binding or phosphorylation. Interestingly, past work has shown a direct interaction between EGFR and another RTK RON (16). Clathrin-coated pits are relatively small structure of ∼100 nm in diameter.

Sequestration of receptors at this scale in subdomains likely contributes to cross-activation and complex and unexpected behaviors though proximity and shared interactions that differ from those seen in biochemical assays.

The PM is a complex and dense environment. Multiple pathways overlap in the same small area of cellular space. Many enzyme scaffolds such as AKAPs have evolved to facilitate these interactions (54). Clathrin appears to act in a similar fashion (55). In this scenario, PM-wide changes in the clathrin system that we previously observed after EGF stimulation likely effects the distribution of other pathways. This behavior adds to the complexity of these growth factor systems. Possible future treatments aimed at combating pathologies that result from the dysregulation of EGFR, FGFR, or LDLR might target these parallel pathways. Indeed, recent drug screens have suggested this is a viable pharmacologic approach for human cancer treatment.

Clathrin participates in cellular roles beyond endocytosis including adhesion, signaling, and cell division (11, 41, 42). Understanding the diversity of these roles is an important frontier in cell biology. Many questions remain. How some cargos and not others are loaded into clathrin is still unclear. If the receptor is eventually endocytosed from these sites at the bottom of the cell or receptors at the top of the cell behave differently is also unknown. In the future, a clearer picture of the mechanisms used by RTKs to orchestrate cellular architecture at the nanoscale will help to identify how to interfere with specific targets to counteract aberrant behaviors in cancer.

## Methods Cell culture

Wild-type HSC3 cells (human oral squamous carcinoma) were obtained from the JCRB Cell Bank (JCRB0623). Previously reported genome-edited HSC-3 cells expressing endogenous EGFR-GFP were generously provided by Dr. Alexander Sorkin (University of Pittsburgh) (21). The cells were cultured at 37 °C with 5% CO_2_ in phenol-free Dulbecco’s modified Eagle’s medium (DMEM) (Thermo-Fisher, Gibco™, 31053028) supplemented with 4.5 g/L glucose, 10% (v/v) fetal bovine serum (Atlanta Biologicals, S10350), 50 mg/mL streptomycin -50 U/mL penicillin (Thermo-Fisher, Gibco™, 15070063), 1% (v/v) Glutamax (Thermo-Fisher, 35050061), and 1 mM sodium pyruvate (Thermo-Fisher, Gibco™, 11360070). The cell lines were used from low-passage frozen stocks and regularly checked for mycoplasma contamination. Transfections were performed by incubating the cells for 4 hours with 500 ng of the specified plasmid(s) and 5 μL of Lipofectamine 3000 (Thermo-Fisher, L3000015) in OptiMEM (Thermo-Fisher, Gibco™, 31985062) according to the manufacturer’s instructions. For experiments, cells were cultured on 25 mm diameter rat tail collagen I-coated coverslips (Neuvitro Corporation, GG-25-1.5-collagen). Experiments were conducted 24-48 hours after transfection with plasmids.

### Plasmids

A complete list of the 57 plasmids used in this study, their construction, and their original sources are included in the supplemental information (SI Appendix, Table 1). All plasmids generated in this study were fully sequenced by Plasmidsaurus.

### Pulse-chase stimulation and drug treatments

Cells were incubated in starvation buffer (DMEM containing 4.5 g/L D-glucose, supplemented with 1% v/v Glutamax and 10 mM HEPES) for 1 h before the pulse-chase assay. Then, cells were pulsed in starvation buffer supplemented with 0.1% w/v bovine serum albumin at 4 °C for 40 min with 50 ng/mL human recombinant EGF (Thermo-Fisher, Gibco™, PHG0311L) to allow ligand bind to the EGFR. We also tested 50 ng/mL human recombinant FGF (Thermo-Fisher, 100-18B), 10 μM noradrenaline (Sigma-Aldrich, A9512-250MG), 1 μM isoproterenol (Sigma-Aldrich, I6504-100MG), 10 μM lysophosphatidic acid (Fisher Scientific, NC9401387). In brief, cells were washed twice with PBS (Thermo-Fisher, Gibco™, 10010023). Synchronized receptor activation and endocytosis were triggered by placing the coverslips in pre-warmed media and incubation at 37 °C for the indicated times. To stop stimulation, cells were washed twice with iced-cold PBS. To block EGFR and FGFR, cells were incubated for 15 min before chase and during pulse with 10 μM gefitinib (Santa Cruz Biotechnology, 184475-35-2), and 1 μM PD-166866 (Selleck Chemicals, S8493), respectively.

### Total Internal Reflection Microscopy (TIRFM)

After pulse-chase stimulation and drugs treatments, cells were fixed with 4 % paraformaldehyde for 20 min at 4 °C and washed 3× with PBS. Cells were imaged on an inverted fluorescent microscope (IX-81, Olympus), equipped with a 100x, 1.45 NA objective (Olympus). Combined green (488 nm) and red (561 nm) lasers (Melles Griot) were controlled with an acousto-optic tunable filter (Andor) and passed through a LF405/488/561/635 dichroic mirror. Emitted light was filtered using a 565 DCXR dichroic mirror on the image splitter (Photometrics), passed through 525Q/50 and 605Q/55 filters and projected onto the chip of an electron-multiplying charge-coupled device (EMCCD) camera. Images were acquired using the Andor IQ2 software. Cells were excited with alternate green and red excitation light, and images in each channel were acquired at 500-ms exposure at 5 Hz. We used a Matlab software previously described (20) to automatically identify clathrin spots in one channel and extract small, square regions centered at the brightest pixel of each object. Matched regions from the same cellular location in the corresponding image were extracted. An equal number of randomly positioned regions were also extracted to test for non-specific colocalization. The mean correlation between thousands clathrin spots and their corresponding image pairs across several cells from independent experiments was calculated using Pearson’s correlation coefficient.

### Statistics

Data were tested for normality and equal variances with Shapiro–Wilk. The statistical tests were chosen as follows: unpaired normally distributed data were tested with a two-tailed *t*-test (in the case of similar variances) or with a two-tailed *t*-test with Welch’s correction (in the case of different variances).

Statistical comparisons between groups were performed using one way ANOVA with Tukey post-test. A *P* value of <0.05 was considered statistically significant. All tests were performed with Origin 2015.

## Acknowledgements

We thank the NHLBI Light Microscopy core for support with fluorescence imaging and instrumentation. We thank members of the Taraska laboratory for discussion and comments on the manuscript. JWT is supported by the Intramural Research Program, National Heart Lung and Blood Institute, National Institutes of Health, Bethesda, Maryland.

## Author contributions

MAAM and JWT designed experiments. MAAM performed experiments. MAAM and JWT analyzed data. MPS performed molecular cloning. MAAM wrote and JWT edited the manuscript and all authors commented on the work. JWT supervised the project.

---

**SI Appendix Fig. 1.**
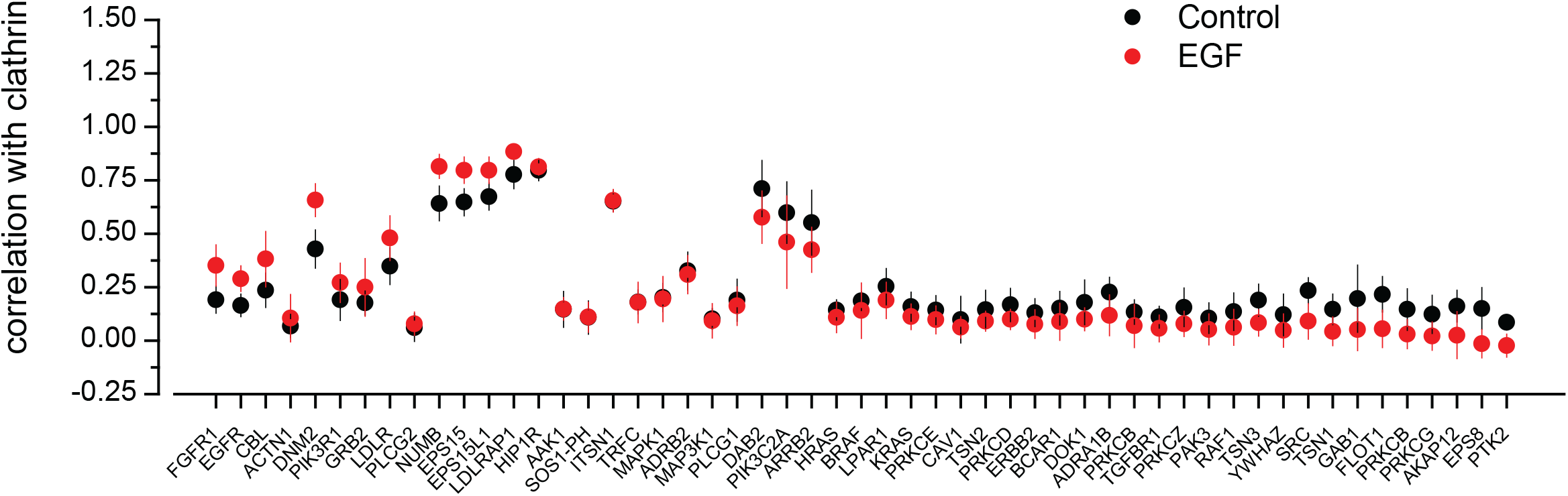
Quantitative measurements of EGF-induced changes to protein correlation with clathrin-coated sites. Automated correlation analysis values of 53 different fluorescently tagged proteins and individual clathrin sites in unstimulated (black) and EGF-stimulated (red) HSC3 cells with a 12 pixel-diameter analysis region. Error is Standard Deviation.

**SI Appendix Table 1.**
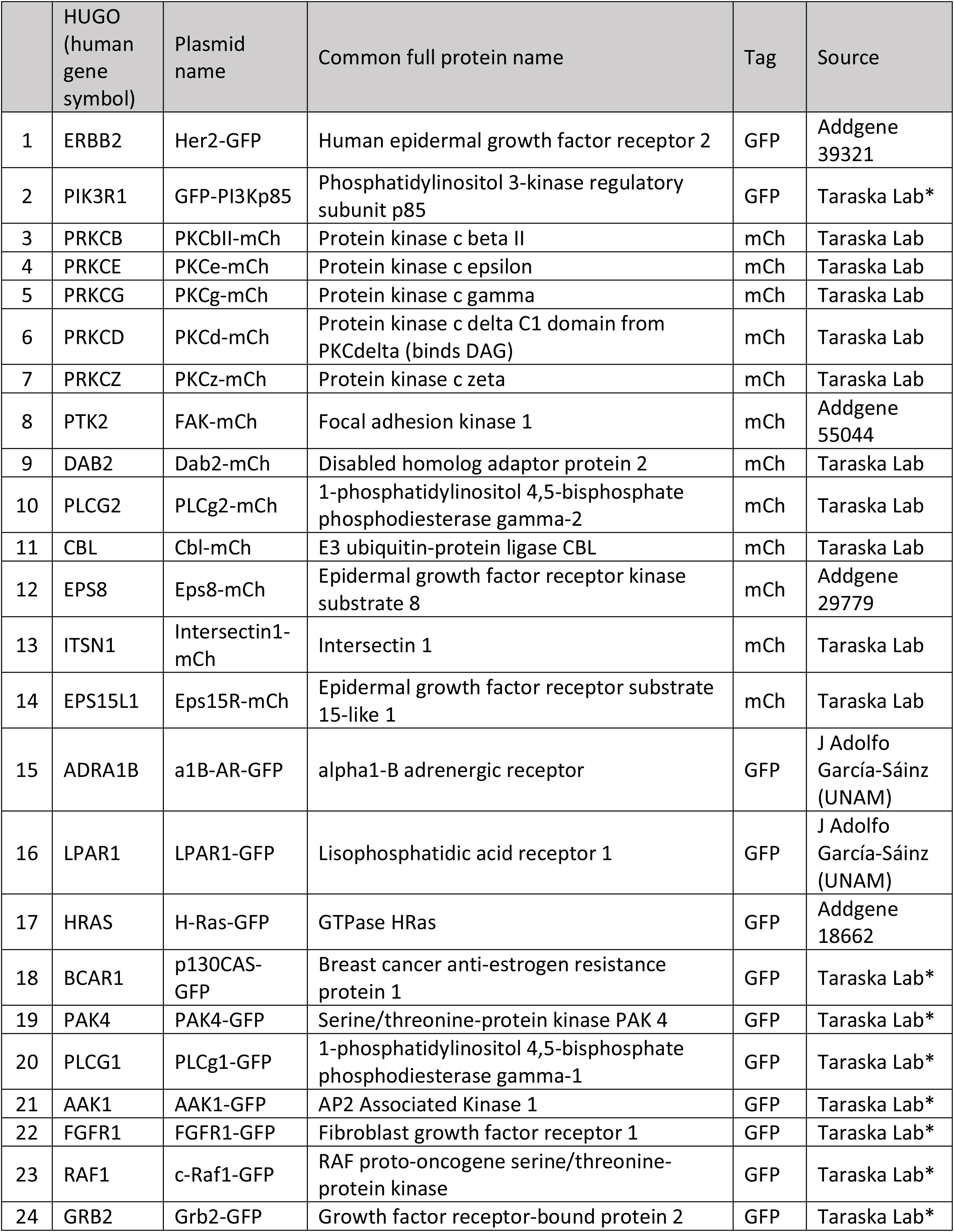
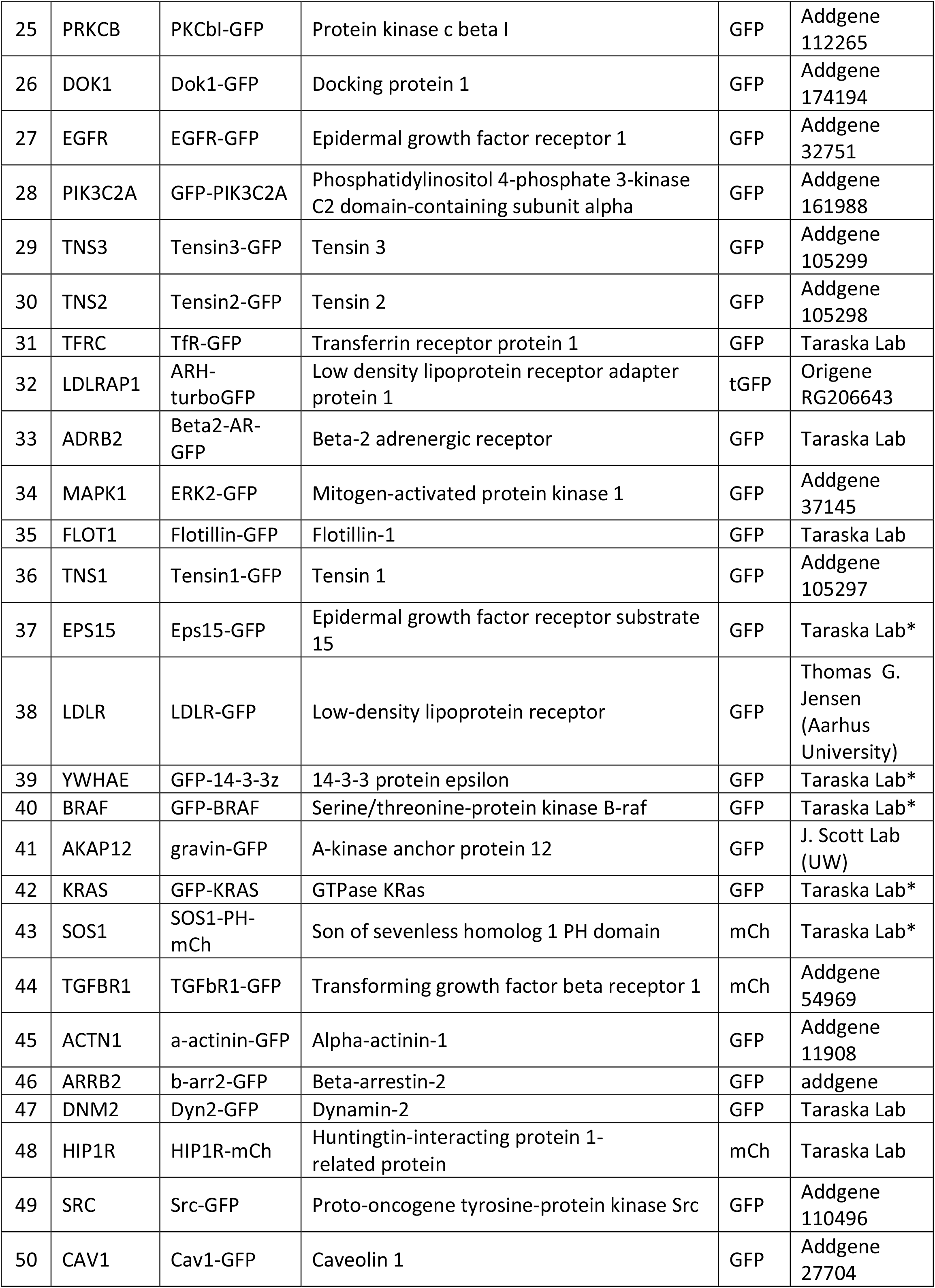
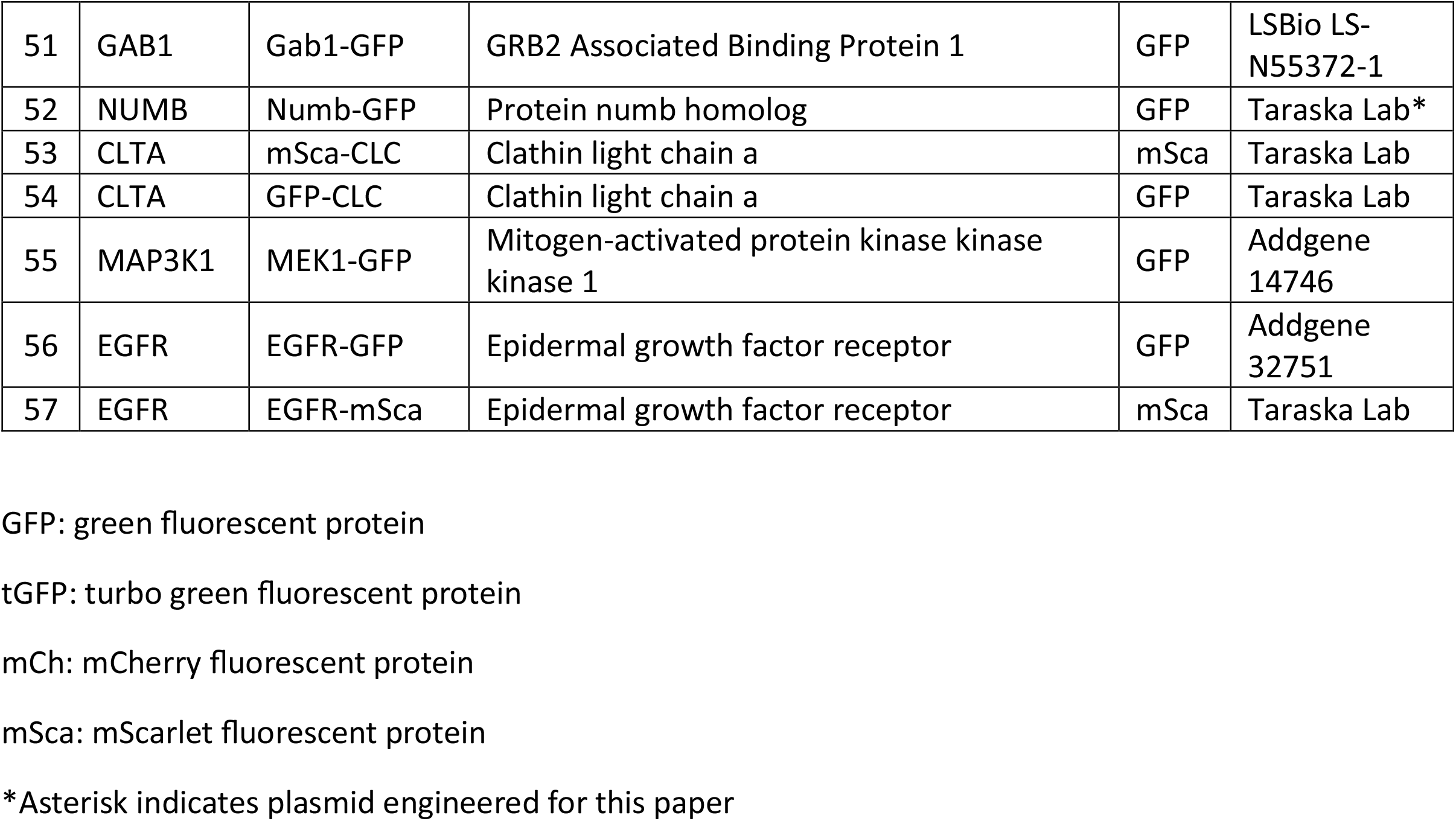
List of plasmids used in the imaging screen.

## References

1. M. A. Lemmon, J. Schlessinger, Cell Signaling by Receptor Tyrosine Kinases. Cell 141, 1117–1134 (2010).

2. S. Sigismund, D. Avanzato, L. Lanzetti, Emerging functions of the EGFR in cancer. Mol Oncol 12, 3–20 (2018).

3. S. Guardiola, M. Varese, M. Sánchez-Navarro, E. Giralt, A Third Shot at EGFR: New Opportunities in Cancer Therapy. Trends Pharmacol Sci 40, 941–955 (2019).

4. H. Ogiso et al., Crystal structure of the complex of human epidermal growth factor and receptor extracellular domains. Cell 110, 775–787 (2002).

5. J. Schlessinger, Ligand-induced, receptor-mediated dimerization and activation of EGF receptor. Cell 110, 669–672 (2002).

6. Y. Kim et al., Temporal Resolution of Autophosphorylation for Normal and Oncogenic Forms of EGFR and Differential Effects of Gefitinib. Biochemistry-Us 51, 5212–5222 (2012).

7. M. P. Verdaguer et al., Time-resolved proximity labeling of protein networks associated with ligand-activated EGFR. Cell Reports 39 (2022).

8. J. Tong, P. Taylor, M. F. Moran, Proteomic analysis of the epidermal growth factor receptor (EGFR) interactome and post-translational modifications associated with receptor endocytosis in response to EGF and stress. Mol Cell Proteomics 13, 1644–1658 (2014).

9. S. L. Schmid, Reciprocal regulation of signaling and endocytosis: Implications for the evolving cancer cell. J Cell Biol 216, 2623–2632 (2017).

10. M. A. Alfonzo-Mendez, K. A. Sochacki, M. P. Strub, J. W. Taraska, Dual clathrin and integrin signaling systems regulate growth factor receptor activation. Nat Commun 13, 905 (2022).

11. D. Leyton-Puig et al., Flat clathrin lattices are dynamic actin-controlled hubs for clathrin-mediated endocytosis and signalling of specific receptors. Nat Commun 8, 16068 (2017).

12. A. Zuidema et al., Mechanisms of integrin alphaVbeta5 clustering in flat clathrin lattices. J Cell Sci 131 (2018).

13. F. Baschieri et al., Frustrated endocytosis controls contractility-independent mechanotransduction at clathrin-coated structures. Nat Commun 9, 3825 (2018).

14. F. A. Sarker, V. G. Prior, S. Bax, G. M. O’Neill, Forcing a growth factor response - tissue-stiffness modulation of integrin signaling and crosstalk with growth factor receptors. Journal of Cell Science 133 (2020).

15. S. P. Kennedy, J. F. Hastings, J. Z. Han, D. R. Croucher, The Under-Appreciated Promiscuity of the Epidermal Growth Factor Receptor Family. Front Cell Dev Biol 4, 88 (2016).

16. C. Franco Nitta et al., EGFR transactivates RON to drive oncogenic crosstalk. Elife 10 (2021).

17. Q. Wu et al., EGFR Inhibition Potentiates FGFR Inhibitor Therapy and Overcomes Resistance in FGFR2 Fusion-Positive Cholangiocarcinoma. Cancer Discov 12, 1378–1395 (2022).

18. S. Sarabipour, K. Hristova, Mechanism of FGF receptor dimerization and activation. Nature Communications 7 (2016).

19. S. Sarabipour, Parallels and Distinctions in FGFR, VEGFR, and EGFR Mechanisms of Transmembrane Signaling. Biochemistry-Us 56, 3159–3173 (2017).

20. B. T. Larson, K. A. Sochacki, J. M. Kindem, J. W. Taraska, Systematic spatial mapping of proteins at exocytic and endocytic structures. Mol Biol Cell 25, 2084–2093 (2014).

21. I. Pinilla-Macua, A. Grassart, U. Duvvuri, S. C. Watkins, A. Sorkin, EGF receptor signaling, phosphorylation, ubiquitylation and endocytosis in tumors in vivo. Elife 6 (2017).

22. A. J. M. Wollman et al., Critical roles for EGFR and EGFR-HER2 clusters in EGF binding of SW620 human carcinoma cells. J R Soc Interface 19, 20220088 (2022).

23. B. Barsi-Rhyne, A. Manglik, M. von Zastrow, Discrete GPCR-triggered endocytic modes enable beta-arrestins to flexibly regulate cell signaling. Elife 11 (2022).

24. M. M. Murph, L. A. Scaccia, L. A. Volpicelli, H. Radhakrishna, Agonist-induced endocytosis of lysophosphatidic acid-coupled LPA1/EDG-2 receptors via a dynamin2- and Rab5-dependent pathway. J Cell Sci 116, 1969–1980 (2003).

25. J. A. Castillo-Badillo et al., alpha1B-adrenergic receptors differentially associate with Rab proteins during homologous and heterologous desensitization. PLoS One 10, e0121165 (2015).

26. T. Scully, N. Kase, E. J. Gallagher, D. LeRoith, Regulation of low-density lipoprotein receptor expression in triple negative breast cancer by EGFR-MAPK signaling. Sci Rep 11, 17927 (2021).

27. D. Leonard et al., Sorting of EGF and transferrin at the plasma membrane and by cargo-specific signaling to EEA1-enriched endosomes. J Cell Sci 121, 3445–3458 (2008).

28. I. Pinilla-Macua, A. Sorkin, Cbl and Cbl-b independently regulate EGFR through distinct receptor interaction modes. Mol Biol Cell 34, ar134 (2023).

29. M. Morimatsu et al., Multiple-state reactions between the epidermal growth factor receptor and Grb2 as observed by using single-molecule analysis. Proc Natl Acad Sci U S A 104, 18013–18018 (2007).

30. K. Abdi et al., EGFR Signaling Termination via Numb Trafficking in Ependymal Progenitors Controls Postnatal Neurogenic Niche Differentiation. Cell Rep 28, 2012–2022 e2014 (2019).

31. M. R. Torrisi et al., Eps15 is recruited to the plasma membrane upon epidermal growth factor receptor activation and localizes to components of the endocytic pathway during receptor internalization. Mol Biol Cell 10, 417–434 (1999).

32. M. Sanford, L. J. Scott, Gefitinib: a review of its use in the treatment of locally advanced/metastatic non-small cell lung cancer. Drugs 69, 2303–2328 (2009).

33. G. Risuleo, M. Ciacciarelli, M. Castelli, G. Galati, The synthetic inhibitor of fibroblast growth factor receptor PD166866 controls negatively the growth of tumor cells in culture. J Exp Clin Cancer Res 28, 151 (2009).

34. M. Lampe, S. Vassilopoulos, C. Merrifield, Clathrin coated pits, plaques and adhesion. J Struct Biol 196, 48–56 (2016).

35. M. von Zastrow, A. Sorkin, Mechanisms for Regulating and Organizing Receptor Signaling by Endocytosis. Annu Rev Biochem 90, 709–737 (2021).

36. R. Irannejad et al., Conformational biosensors reveal GPCR signalling from endosomes. Nature 495, 534–538 (2013).

37. J. G. Lock et al., Clathrin-containing adhesion complexes. J Cell Biol 218, 2086–2095 (2019).

38. W. F. Zeno et al., Clathrin senses membrane curvature. Biophys J 120, 818–828 (2021).

39. M. Kaksonen, A. Roux, Mechanisms of clathrin-mediated endocytosis. Nat Rev Mol Cell Biol 19, 313–326 (2018).

40. M. Lehmann et al., Nanoscale coupling of endocytic pit growth and stability. Sci Adv 5, eaax5775 (2019).

41. J. G. Lock et al., Reticular adhesions are a distinct class of cell-matrix adhesions that mediate attachment during mitosis. Nat Cell Biol 20, 1290–1302 (2018).

42. L. Hakanpaa et al., Reticular adhesions are assembled at flat clathrin lattices and opposed by active integrin alpha5beta1. J Cell Biol 222 (2023).

43. F. Lukas et al., Canonical and non-canonical integrin-based adhesions dynamically interconvert. Nat Commun 15, 2093 (2024).

44. A. Franck et al., Clathrin plaques and associated actin anchor intermediate filaments in skeletal muscle. Mol Biol Cell 30, 579–590 (2019).

45. J. H. Bae et al., Asymmetric receptor contact is required for tyrosine autophosphorylation of fibroblast growth factor receptor in living cells. Proc Natl Acad Sci U S A 107, 2866–2871 (2010).

46. E. Salazar-Cavazos et al., Multisite EGFR phosphorylation is regulated by adaptor protein abundances and dimer lifetimes. Mol Biol Cell 31, 695–708 (2020).

47. S. K. Basu, J. L. Goldstein, R. G. Anderson, M. S. Brown, Monensin interrupts the recycling of low density lipoprotein receptors in human fibroblasts. Cell 24, 493–502 (1981).

48. M. S. Brown, R. G. Anderson, J. L. Goldstein, Recycling receptors: the round-trip itinerary of migrant membrane proteins. Cell 32, 663–667 (1983).

49. S. Amos et al., Protein kinase C-alpha-mediated regulation of low-density lipoprotein receptor related protein and urokinase increases astrocytoma invasion. Cancer Res 67, 10241–10251 (2007).

50. D. R. Hummel, M. Kaksonen, Spatio-temporal regulation of endocytic protein assembly by SH3 domains in yeast. Mol Biol Cell 34, ar19 (2023).

51. U. Dionne, L. J. Percival, F. J. M. Chartier, C. R. Landry, N. Bisson, SRC homology 3 domains: multifaceted binding modules. Trends Biochem Sci 47, 772–784 (2022).

52. K. J. Day et al., Liquid-like protein interactions catalyse assembly of endocytic vesicles. Nat Cell Biol 23, 366–376 (2021).

53. A. Patwardhan, N. Cheng, J. Trejo, Post-Translational Modifications of G Protein-Coupled Receptors Control Cellular Signaling Dynamics in Space and Time. Pharmacol Rev 73, 120–151 (2021).

54. M. H. Omar, J. D. Scott, AKAP Signaling Islands: Venues for Precision Pharmacology. Trends Pharmacol Sci 41, 933–946 (2020).

55. S. Vassilopoulos et al., Actin scaffolding by clathrin heavy chain is required for skeletal muscle sarcomere organization. J Cell Biol 205, 377–393 (2014).

